# Virtual cortical resection reveals push-pull network control preceding seizure evolution

**DOI:** 10.1101/055566

**Authors:** Ankit N. Khambhati, Kathryn A. Davis, Timothy H. Lucas, Brian Litt, Danielle S. Bassett

## Abstract

For ≈ 20 million people with drug-resistant epilepsy, recurring, spontaneous seizures have a devastating impact on daily life. The efficacy of surgical treatment for controlling seizures is hindered by a poor understanding of how some seizures spread to and synchronize surrounding tissue while others remain focal. To pinpoint network regions that regulate seizure evolution, we present a novel method to assess changes in synchronizability in response to virtually lesioning cortical areas in a validated computational network model. In human patients implanted with electrocorticographic sensors, we apply our virtual cortical resection technique to time-varying functional networks and identify control regions that synchronize or desynchronize cortical areas using an antagonistic push-pull control scheme to raise or lower synchronizability. Our results suggest that synchronizability before seizures predicts seizure evolution: in focal seizures, the strongest controllers are located outside seizure-generating areas. These methods,while applied here to epilepsy, are generalizable to other brain networks, and have wide applicability in isolating and mapping functional drivers of brain dynamics in health and disease.

## 1 Introduction

Functional architecture of the epileptic neocortex has been studied extensively to better identify optimal targets for surgical resection and, more recently, the optimal location for focal ablation or implantable devices [1, 2, 3]. The prospect of patient-centric algorithms that modulate brain state to abort seizures is exciting to clinicians and researchers alike [4, 5, 6]. However, the best targets for chronic devices remain elusive, partly because functional brain networks, including epileptic networks, reorganize dynamically [7, 8, 9, 10, 11]. Such reorganization appears to follow a specific progression through network states unique to the patient,s seizures [12, 10, 11]. The mechanisms that drive seizures through network states can inform neural control paradigms that aim to stop or contain propagation of seizure activity. Such a capability is vital, clinically, because epileptogenic regions cause symptoms not only through their own dysfunction, but also through their ability to recruit and disrupt healthy brain tissue [13]. Understanding and translating network mechanisms of seizure evolution to identify targets for therapy requires further intellectual dissection of functional epileptic network architecture.

Conventional thinking divides epileptic brain into clinically-defined regions where seizures presumably originate [14] and surrounding regions in which seizures do not originate. Recent models describe connectivity between seizure-onset and surrounding cortical regions in the framework of a broader dysfunctional *epileptic network*, where network nodes are neural populations measured by intracranial sensors and network connections are statistical relationships between neural activation patterns [15, 16, 17, 18, 10, 11] (**Fig. 1a**). For example, partial seizures that begin in the seizure-onset zone can evolve, spreading spatially as they modulate in dominant frequency via local connections to the surrounding tissue, implicating a distributed epileptic network [19, 15, 16, 20, 11]. In the extreme case, these seizures can generalize and eventualy encompass the entire brain.

Given the distributed nature of epileptic activity, it is critical to isolate underlying propagation mechanisms. Leading hypotheses suggest that either (i) seizure evolution is driven by strong, synchronizing activity from the seizure-generating network impinging outward on the surrounding tissue [21, 16, 22, 23], or (ii) seizure evolution is caused by a diminished ability of the surrounding tissue to regulate, or contain, abnormal activity [15, 24]. While little evidence exists to determine which of these hypotheses accurately reflect seizure dynamics, both mechanisms can be succinctly summarized as abnormalities of synchronizability, a description of how easily neural processes, such as rhythmic activity, can diffuse through a network.

Theoretical work in the fields of physics and engineering demonstrates that diffusion of dynamics through the network can be regulated through a *push-pull control* mechanism, where desynchronizing and synchronizing nodes operate antagonistically in a “tug-of-war”. When synchronizing nodes exert greater push than desynchronizing nodes, synchronizability increases and dynamic processes may diffuse through the network more easily [25] (****Fig. 1b****). Such mechanisms are particularly successful in heterogeneous networks like the brain, where some nodes are sparsely connected and other nodes are densely connected [26]. Does the brain utilize such a control mechanism for seizure regulation? And if so, what regions of the brain affect this control?

To address these questions, we present a novel method we call *virtual cortical resection,* which offers a statistically robust means to pinpoint putative control nodes in the epileptic network that may regulate seizure dynamics, based on the network,s response to virtual lesioning [26, 27]. We use this method to test the hypothesis that the epileptic network contains a native regulatory system (**Fig. 1c**) whose connectivity to the seizure-generating area accounts for differential seizure dynamics, including (i) the constrained dynamics observed in partial seizures that remain focal (**Fig. 1d**), and (ii) the unconstrained dynamics observed in partial seizures that generalize to surrounding tissue (**Fig. 1e**).

More specifically, using electrocorticography recorded from 10 patients diagnosed with drug-resistant neocortical epilepsy undergoing routine pre-surgical evaluation, we constructed time-evolving functional networks across *events*, each of which included a seizure epoch preceded by a pre-seizure epoch. The seizure epoch spanned the period between the clinically-marked earliest electrographic change [28] and seizure termination, while the pre-seizure epoch was identical in duration to the seizure and ended immediately prior to the earliest electrographic change. In each epoch, we divided the ECoG signal into 1s non-overlapping time-windows and estimated functional connectivity in high-*γ* (95–105 Hz) and low-*γ* (30–40 Hz) frequency bands using multitaper coherence estimation (see *Methods*). We implemented virtual cortical resection on this dynamic epileptic network by independently removing electrode sites from the network model. This was done to assess the synchronizability of (i) the distributed epileptic network in partial seizures that *generalize* to surrounding tissue, *versus* (ii) the focal epileptic network in seizures that *do not generalize* to surrounding tissue. By removing electrode sites from the network model, we were able to probe the importance of brain regions, in their presence and absence, to seizure generation and propagation.

**Figure 1:**
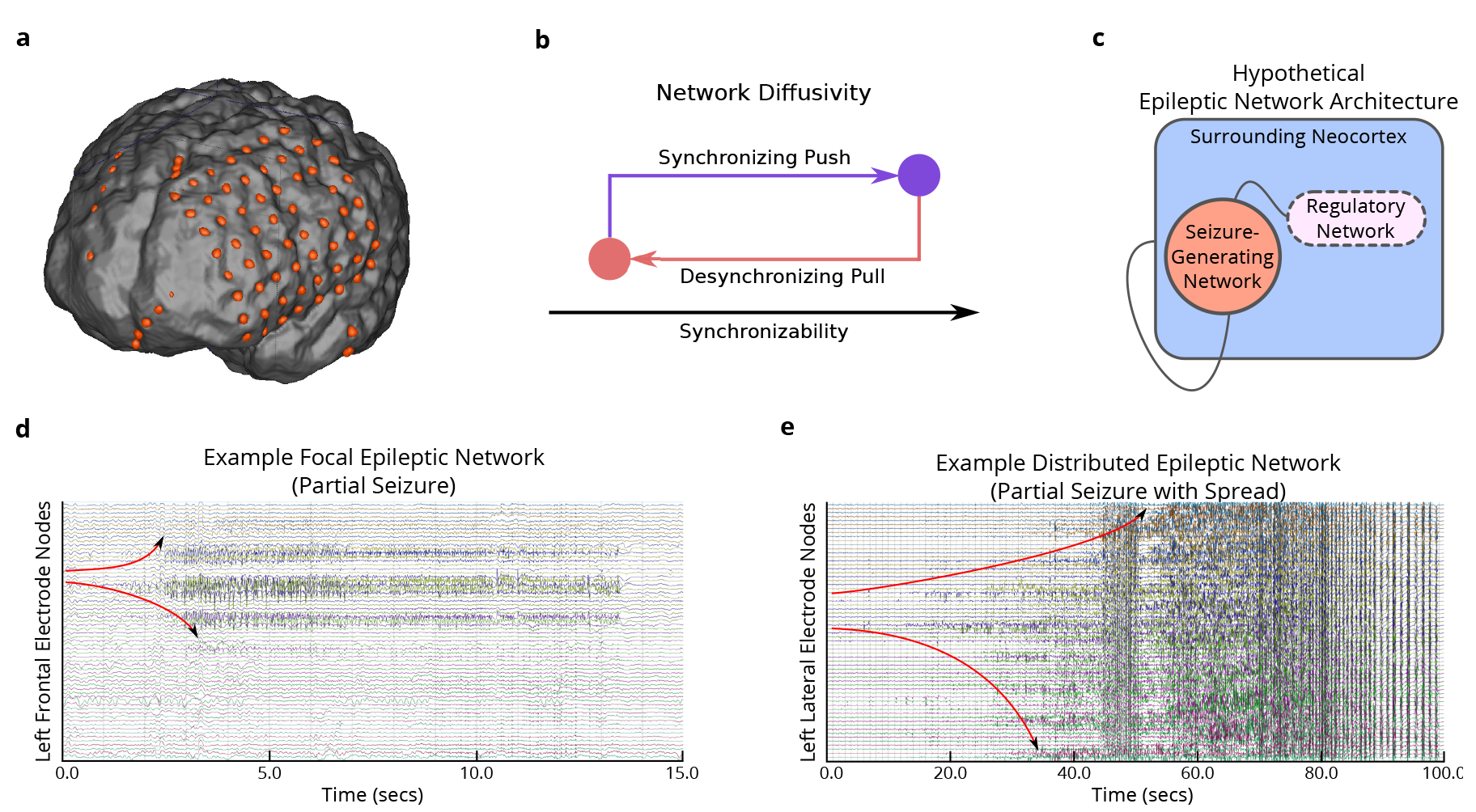
Hypothesized Mechanism of Seizure Regulation. (**a**) We created functional networks based on electrophysiology from patients with drug-resistant neocortical epilepsy implanted with intracranial electrodes. Each sensor is represented as a network node, and weighted functional connectivity between sensors, interpreted as degree of coherence, is represented as a network connection. (**b**) Rope-stretching diagram demonstrating push-pull control, where opposing synchronizing and desynchronizing forces (nodes) shifts overall network synchronizability. (**c**) Schematic of the epileptic network composed of a *seizure-generating system* and a hypothesized *regulatory system* that controls the spread of pathologic seizure activity. (**d**) Example partial seizure that remains focal: the seizure begins at a single node and evolves to and persists within a focal area. (**e**) Example partial seizure that generalizes to surrounding tissue: the seizure begins at two nodes and evolves to the broader network. We hypothesize that these two types of dynamics are determined by differences in the regulatory system.

## 2 Results

### 2.1 Network Homogeneity Improves Synchronizability

We first asked the question, “How easily do seizures diffuse through distributed and focal epileptic networks?” We hypothesized that spread of seizures can be quantified by the network synchronizability, or potential for the network to synchronize due to seizures. To quantify network synchronizability, we estimated the time-varying Laplacian matrix **L**(*t*) whose entries *l_ij_*(*t*) quantify how easily information can diffuse between nodes *i* and *j* (see *Methods*). Using the Laplacian matrix, we computed the synchronizability 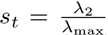, where λ_2_ and λ_max_ are the second-smallest eigenvalue and the largest eigenvalue, respectively, of **L**(*t*) (see [29] and **Supplementary Note**). Intuitively, greater network synchronizability implies greater ease for neural populations to synchronize their dynamics - such as during seizures. We observed significantly greater synchronizability in the distributed epileptic network than in the focal epileptic network during the preseizure epoch, suggesting that high-γ networks have a greater potential to synchronize prior to seizures that spread than prior to seizures that do not spread (**Fig. 2a**). In contrast, we observed synchronizability in low-γ networks effectively captured spread through the distributed epileptic networks after seizure-onset (see *Supplemental Information:* **Fig. S1a**).

**Figure 2:**
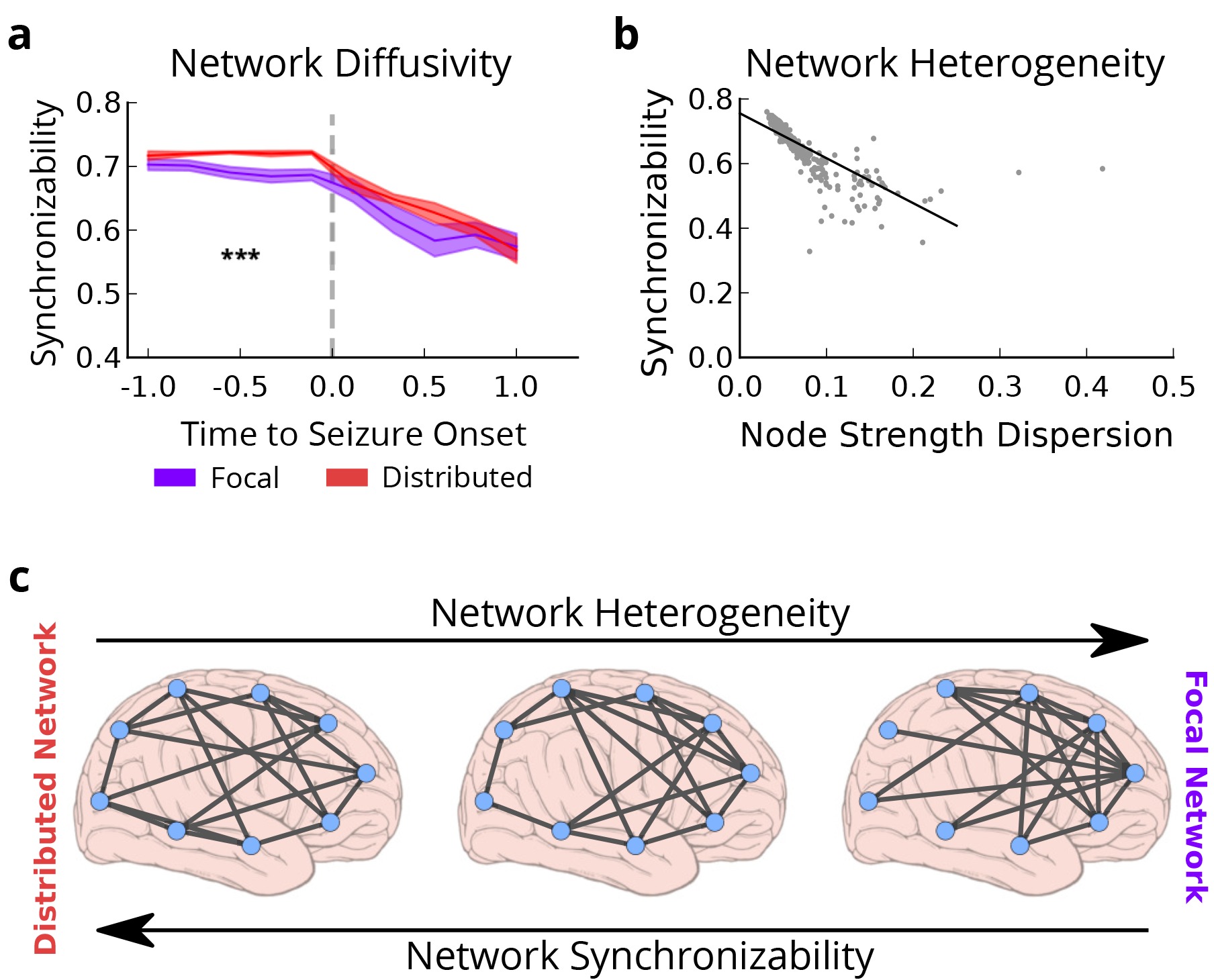
Differential Pre-Seizure Synchronizability Predicts Seizure Spread. (**a**) Time-dependent synchronizability captures the potential for seizure spread through high-γ functional networks. Distributed epileptic networks describe seizures with secondary generalization (N = 16), focal epileptic networks describe seizures without secondary generalization (N = 18). Analyzed epileptic events spanned the clinically-defined seizure and period of time equal in duration to the seizure, immediately preceding seizure-onset. Events were time-normalized with each pre-seizure and seizure period divided into 5 equally-spaced time bins (10 bins per event). Synchronizability was averaged within each bin. Synchronizability was significantly greater in distributed epileptic networks than in focal epileptic networks prior to seizure onset (functional data analysis, *p*_pre-seizure_ = 1.7 × 10^−4^, *p*_seizure_ = 3.1 × 10^−1^). Thick lines represent mean, shaded area represents standard error around mean. *P*-values are obtained via the statistical technique known as functional data analysis (FDA) where event labels (two seizure types) were permuted uniformly at random (see Methods): ^***^*p* < 0.001. (**b**) Relationship between synchronizability and dispersion of node strengths in high-γ functional networks across the population of distributed and focal epileptic network events. Each point represents average synchronizability and dispersion of average node strengths from a single time bin (N = 340). Greater synchronizability was strongly related to greater network heterogeneity, or lower node strength dispersion (Pearson correlation coefficient; *r* = −0.811, *p* < 1 × 10^−16^). (**c**) Schematic demonstrating that distributed epileptic networks have greater synchronizability and more homogeneous topology than focal epileptic networks. Seizures may spread more easily in distributed epileptic networks due to more homogeneous connectivity between network nodes.

Next, based on theoretical work in physics and engineering, we asked if network synchronizability, or predisposition to seizure spread, might be explained by heterogeneity in network topology. That is, “Does heterogeneity in node strength weaken the network,s ability to synchronize?” To measure heterogeneity, we computed a non-parametric, normalized measure of node strength dispersion *d(t)* for each time window *t* (see *Methods*). More heterogeneous network topologies would incur greater node strength dispersion, suggesting nodes might either be highly connected or highly isolated (whereas lower node strength dispersion suggests nodes are more evenly connected in the network). Our results demonstrated a significant linear relationship between synchronizability and node strength dispersion (Pearson correlation; *r* = −0.811, *p* < 1 × 10^−16^), where greater heterogeneity in node strength lead to lower synchronizability (**Fig. 2b**). The correlation between synchronizability and node strength dispersion was also observed in low-*γ* functional networks (see Supplement), suggesting a fundamental and robust relationship between these topological measures.

More generally, our results suggest that seizure spread in the distributed epileptic network may result from a vulnerability to synchronize easily, a vulnerability that is not present in partial seizures that do not generalize to surrounding tissue. Furthermore, the heightened synchronizability of distributed epileptic networks may be driven by homogenous distributions of connectivity amongst network nodes.

### 2.2 Network Controllers of Synchronizability

How might network nodes regulate levels of synchronizability? Do a subset of nodes act as key controllers, or do all nodes contribute equally? To answer this question, we developed a novel method to assess the influence of a node on synchronizability. We define the *control centrality c_i_* of node *i* to be the fractional change in synchronizability following removal of node *i* from the network (**Fig. 3a**): 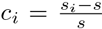where *s* is the original synchronizability and *s_i_* is the synchronizability after node removal. The magnitude of *c_i_* can be interpreted as the overall strength of the node as a controller of synchronizability. If *c_i_* is positive, then synchronizability increases upon node removal, and the node is said to be a *desynchronizing node.* If *c_i_* is negative, then synchronizability decreases upon node removal, and the node is said to be a *synchronizing node.* As illustrated in **Fig. 3a**, both desynchronizing and synchronizing network controllers are characteristic of heterogeneous networks, and tend to be located in the network periphery and network core, respectively.

**Figure 3:**
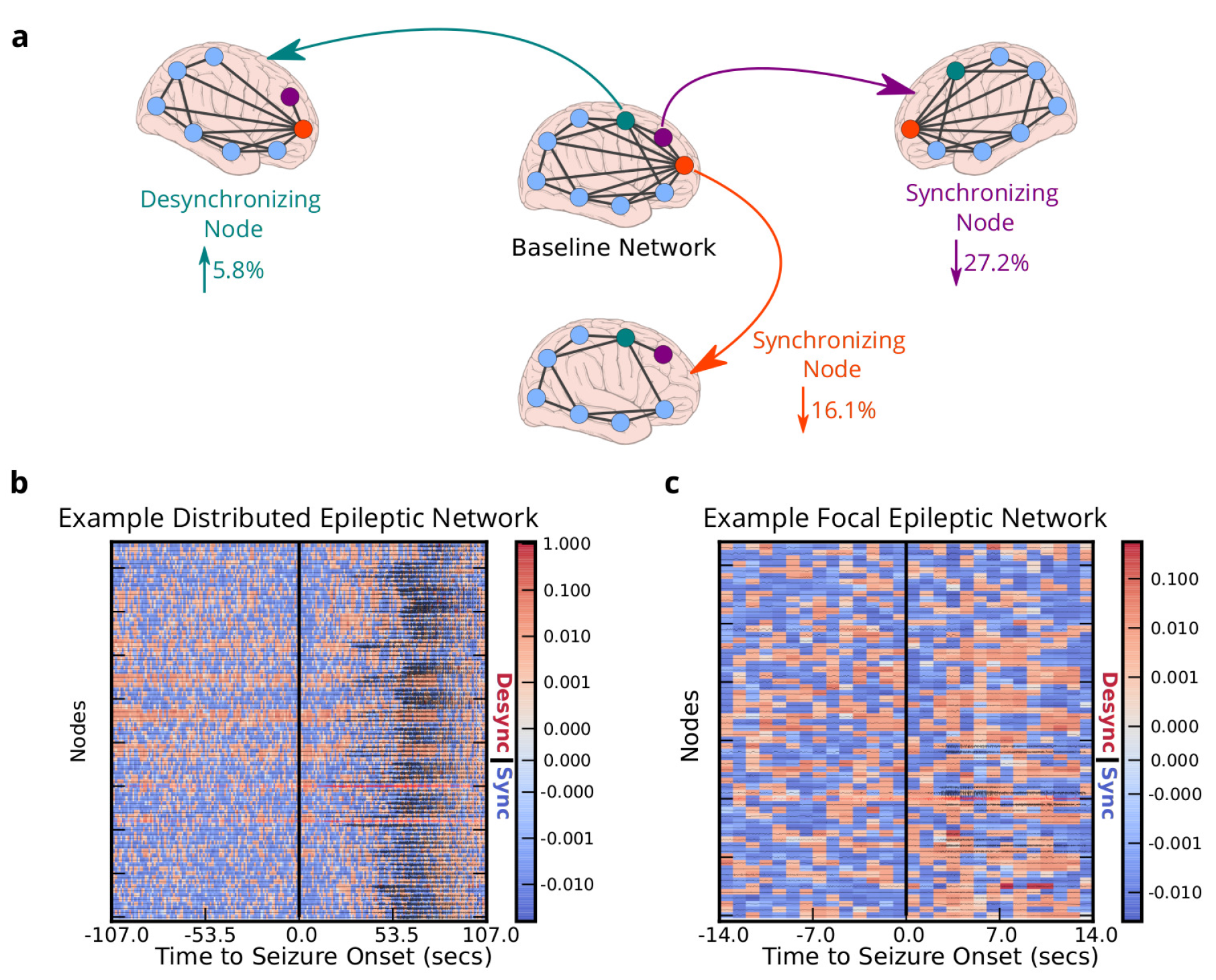
Virtual Cortical Resection Localizes Network Controllers. (**a**) Effect of node removal on network synchronizability (control centrality) in a toy network. Highlighted node removals resulting in increased synchronizability (desynchronizing node; green) or decreased synchronizability (synchronizing nodes; purple and orange). The strongest desynchronizing node increased synchronizability by 5.8% and was present in the network periphery, while the strongest synchronizing nodes decreased synchronizability by 27.2% and 16.1% and were located in the network core. (**c**) Virtual cortical resection applied to example distributed (and (**d**) focal) high-γ epileptic network events. Control centrality of nodes is indicated by color in each time window, and ECoG signal is overlayed and normalized by maximum amplitude (red signals are clinically-defined seizure onset nodes). Control centrality values are positive/red (negative/blue) for desynchronizing (synchronizing) nodes.

We used control centrality to assess the presence of desynchronizing and synchronizing controllers in the epileptic network, and to define their putative role in regulating synchronizability, a hallmark of seizure spread. In example distributed (**Fig. 3b**) and focal (**Fig. 3c**) epileptic networks, we observed clusters of synchronizing and desynchronizing control nodes that differed in their temporal dynamics, as well as in their spatial distribution. How are synchronizing and desynchronizing regions distributed in the epileptic network? Furthermore, can the spatial and temporal distributions of control nodes differentially explain seizure spread?

### 2.3 Regulatory System Controls Seizure Dynamics

To address these questions, we explored regional control of seizure spread in high-γ functional networks (see *Supplemental Information* for analysis of low-γ functional networks). To assign network regions, a team of neurologists successfully identified the sensors on the seizure onset zone (SOZ) based on visual inspection of the intracranial recordings. Sensors within the SOZ were grouped as the seizure onset region, while sensors outside the SOZ were labelled as the surrounding region. Within each region, we computed control centrality for the 10% strongest desynchronizing and synchronizing nodes during pre-seizure and seizure epochs.

First, we compared control centrality of nodes within the seizure-onset region during the pre-seizure epoch (**Fig. 4a**). In focal epileptic networks, control centrality was significantly greater among desynchronizing nodes than synchronizing nodes (Wilcoxon rank-sum; *z* = 2.53, *p* = 0.011). In distributed epileptic networks, we observed no significant difference in control centrality between desynchronizing and synchronizing nodes (Wilcoxon rank-sum; *z* = 1.58, *p* = 0.113). However, SOZ nodes of the distributed network were significantly more synchronizing than similar nodes of the focal network (Wilcoxon rank-sum; *z* = 2.03, *p* = 0.041). No significant difference in desynchronizing strength of SOZ nodes was found between focal and distributed networks (Wilcoxon rank-sum; *z* = 0.03, *p* = 0.972). We then compared control centrality of nodes within the surrounding region during the pre-seizure epoch (**Fig. 4b**). In focal epileptic networks, control centrality was significantly greater among desynchronizing nodes than synchronizing nodes (Wilcoxon rank-sum; *z* = 2.27, *p* = 0.023). In distributed epileptic networks, no significant difference in control centrality between desynchronizing and synchronizing nodes was found (Wilcoxon rank-sum; *z* = 1.54, *p* = 0.122).

**Figure 4:**
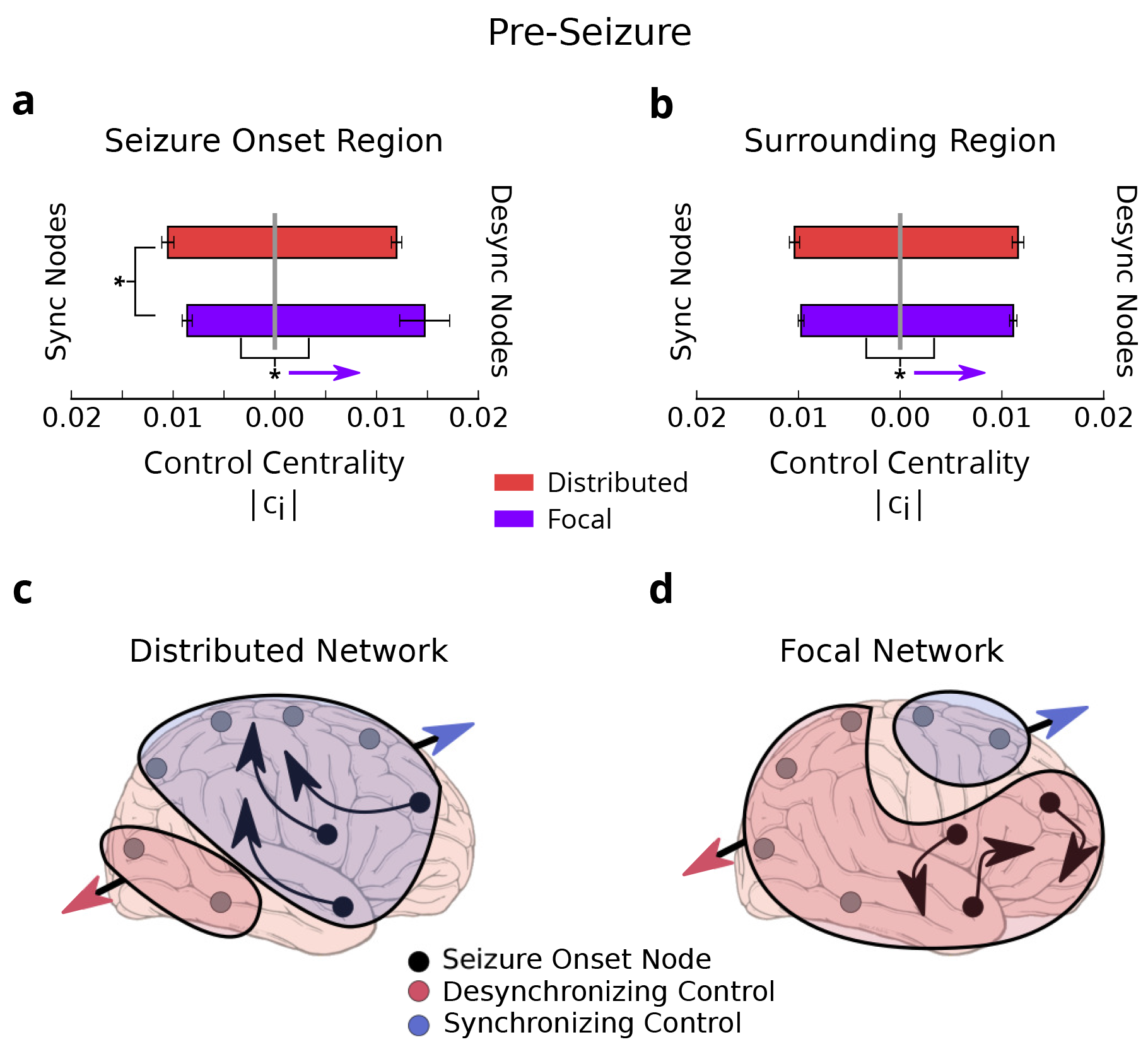
Regional Control Centrality Differentiates Seizure Type in Pre-Seizure Epoch. (**a**) Distribution of control centrality in the 10% strongest synchronizing and desynchronizing nodes within the seizure onset region of focal (N = 18) and distributed (N = 16) events. Focal networks have stronger desynchronizing nodes than synchronizing nodes. Distributed networks have stronger synchronizing nodes than focal networks. ^*^*p* < 0.05. (**b**) Distribution of control centrality in 10% strongest synchronizing and desynchronizing nodes within the surrounding region of focal (N = 18) and distributed (N = 16) events. Focal networks have stronger desynchronizing nodes than synchronizing nodes. (**c**) Schematic of strong synchronizing control in seizure onset and surrounding region of distributed networks that may ease seizure spread. (**d**) Schematic of strong desynchronizing control in seizure onset and surrounding region of focal networks that may regulate seizure spread.

These findings suggest that (i) SOZ nodes are more synchronizing in distributed networks, (ii) SOZ nodes are more desynchronizing in focal networks, and (iii) surrounding regions are more strongly desynchronizing than synchronizing in focal networks (**Fig. 4c-d**). Importantly, our observation of lower synchronizability in the pre-seizure epoch of focal networks coincided with strong desynchronizing control from the SOZ and surrounding region.

Does control centrality of different brain regions distinguish spread in focal and distributed networks during seizures? First, we compared control centrality within the seizure-onset region during the seizure epoch. In focal epileptic networks, control centrality was significantly greater among desynchronizing nodes than synchronizing nodes (Wilcoxon rank-sum; *z* = 2.34, *p* = 0.019). In distributed epileptic networks, we observed no significant difference in control centrality between desynchronizing and synchronizing nodes (Wilcoxon rank-sum; *z* = 1.17, *p* = 0.243). However, SOZ nodes of the distributed network were significantly more synchronizing than similar nodes of the focal network (Wilcoxon rank-sum; *z* = 2.76, *p* = 0.005). No significant difference in control centrality of desynchronizing SOZ nodes was found between focal and distributed networks (Wilcoxon rank-sum; *z* = 0.66, *p* = 0.512). We then compared control centrality of nodes within the surrounding region during the seizure epoch (**Fig. 5b**). We observed significantly stronger synchronizing nodes in distributed networks compared to focal networks (Wilcoxon rank-sum; *z* = 2. 76, *p* = 0.038). We also observed significantly stronger desynchronizing nodes in distributed networks compared to focal networks (Wilcoxon rank-sum; *z* = 2.31, *p* = 0.021).

**Figure 5:**
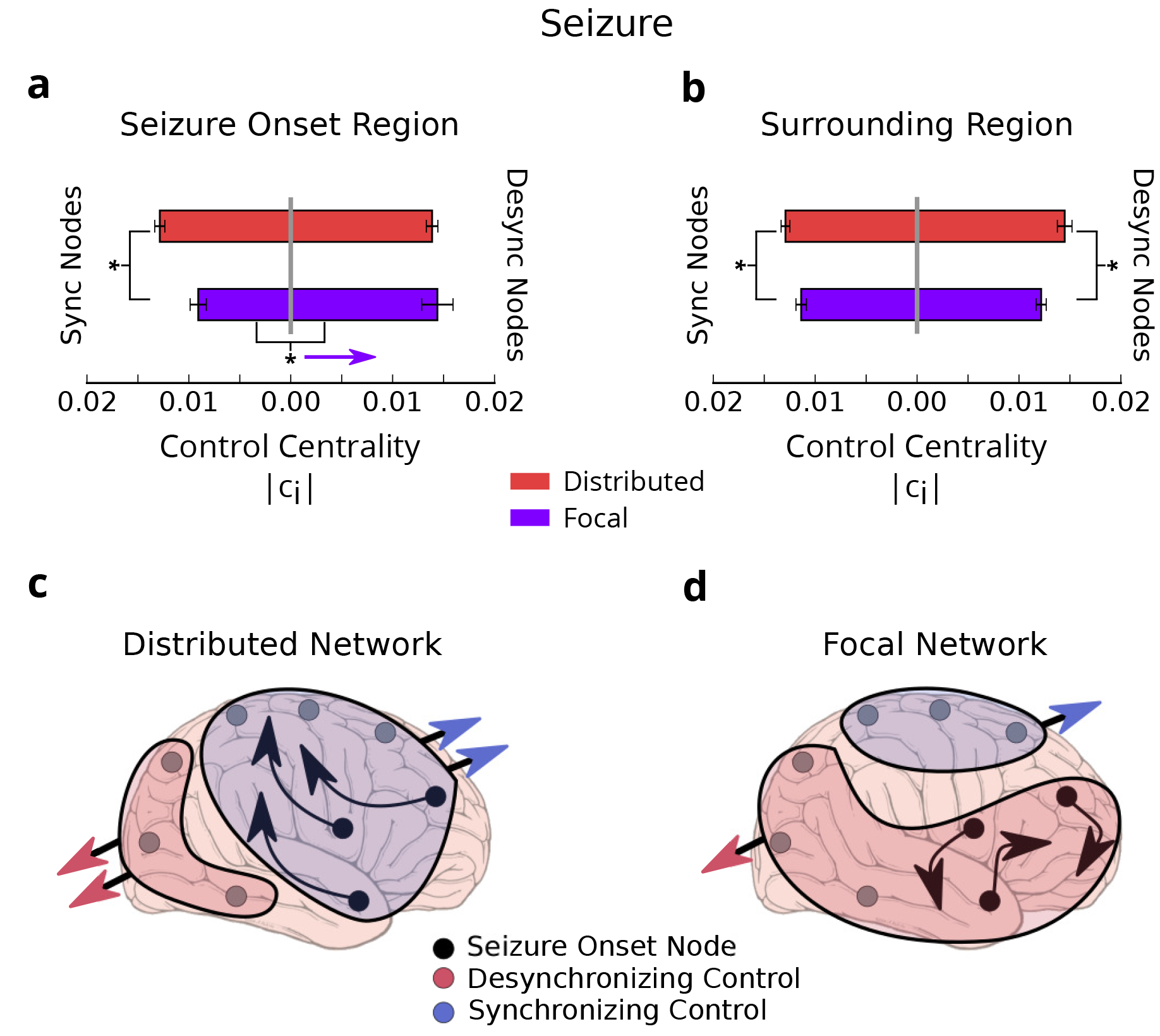
Regional Control Centrality Differentiates Seizure Type in Seizure Epoch. (**a**) Distribution of control centrality in 10% strongest synchronizing and desynchronizing nodes within the seizure onset region of focal (N = 18) and distributed (N = 16) events. Focal networks have stronger desynchronizing nodes than synchronizing nodes. Distributed networks have stronger synchronizing nodes than focal networks. ^*^*p* < 0.05. (**b**) Distribution of control centrality in 10% strongest synchronizing and desynchronizing nodes within the surrounding region of focal (N = 18) and distributed (N = 16) events. Distributed networks have stronger synchronizing and desynchronizing nodes than focal networks. (**c**) Schematic of strong synchronizing control in seizure onset and surrounding region of distributed networks that may ease seizure spread. (**d**) Schematic of strong desynchronizing control in seizure onset and surrounding region of focal networks that may regulate seizure spread.

These findings suggest that during seizures, control centrality of SOZ nodes maintain their respective roles as synchronizing controllers in distributed networks and desynchronizing controllers in focal networks (**Fig. 5c-d**). However, critical changes in the surrounding region occurs after seizures begin: (i) all controller types become stronger, (ii) distributed networks exhibit stronger synchronizing and desynchronizing control than focal networks, and (iii) strong desynchronizing control in focal networks is countered by strong synchronizing control.

Overall, virtual resection of network nodes revealed putative controllers of synchronizability during seizures. We observed strong desynchronizing control within seizure onset and surrounding regions of focal networks that may play a mechanistic role in the regulation of seizure spread by lowering network synchronizability prior to the start of seizures. In contrast, synchronizing control effectively counter-balanced desynchronizing control in distributed networks that demonstrated seizure spread.

## 3 Discussion

In this work we asked, “Is there a network-level control mechanism that regulates seizure evolution?” To answer this question, we designed and applied a novel computational tool - *virtual cortical resection* - to predict network response to removing regions in the epileptic network. We showed network topology preemptively facilitates seizure spread by regulating the synchronizability of epileptic activity. Specifically, synchronizing and desynchronizing network regions competitively modulate network synchronizability and constrain seizure spread beyond seizure-generating regions, effectively forming a push-pull control system. Our results not only significantly extend our understanding of the putative mechanisms of seizure evolution, but also provide a model ripe for further investigation in other brain network disorders and cognition.

### 3.1 Synchronization of the Epileptic Network

Epilepsy researchers have long desired an answer to the question of”Which brain regions drive seizure generation and evolution?” The literature frequently describes the epileptic brain as one that displays an imbalance of excitatory and inhibitory neural populations, leading to instabilities in synchronization that drive seizure dynamics. Clinical observation has informed the development of a dictionary of electrophysiological biomarkers believed to manifest as a result of dysfunctional imbalances in neural populations. However, the variability among epileptologists,s descriptions of epileptic events often leads to poor performance of algorithms that seek to mimic the clinical characterization. In addition, more recent work looking at the cellular level indicates that seizure generation is much more complex and involves interctions between a much wider population of neuronal subtypes both within and surrounding seizure-generating regions [30, 31]

In this work, we studied synchronizability: an objective measure of the ability for neural populations to synchronize in the network. We used this measure to describe mechanistic differences between seizures that either remain focal or that spread throughout the network. Prior work has explored differences in seizure semiology to describe primary and secondary zones of dysfunction [14, 15]. Others have identified strong, tightly connected network hubs localized in areas outside o the seizure-generating region that indicate a wider extent of network damage [32, 33, 9]. Our results support the view that tissue surrounding the seizure-generating area displays abnormalities that support seizure evolution. Specifically, we observe the presence of putative control nodes within a broader heterogeneous network that may serve to oppose seizure spread by limiting synchronizability of healthy activity states.

Our findings supporting the existence of a control system that manages the degree of synchrony in the cortical network may have important neurobiological implications. They raise questions such as,”What is the neuroanatomical substrate for network nodes that drive or contain seizures?” Might there be direct anatomical dysfunction such as loss of inhibitory inter-neurons, aberrant fiber-sprouting or changes in local gap junctions or ion channel expression that correlate with desynchronizing or synchronizing functional regions? Relating correlates of dysfunction from node resection and electrophysiologic studies to underlying neuroanatomy in applications of targeted drug-delivery remains a promising area of epilepsy research.

### 3.2 Push-Pull Control Titrates Network Synchronizability

Controllability of brain networks is a burgeoning area of network neuroscience, particularly in the study of large-scale brain areas and the distributed circuits that they constitute [7, 34, 35]. However, an understanding of the principles of brain network control may have even greater impact in the context of meso-scale brain networks, where local neural populations frequently switch between a wide variety of normal and abnormal rhythmic neural processes. Using virtual cortical resection, we observed the presence of specific nodes whose placement in the wider network suggests their critical role in controlling synchronization and desynchronization in seizure dynamics. These key areas display antithetical potential for controlling activity dynamics, and therefore we speculate that they may employ an antagonistic, push-pull control mechanism similar to that described in theoretical work in other systems [25]. Mechanistically, synchronizing controllers theoretically pull the network towards a particular synchronous state, and, conversely, desynchronizing controllers push the network away from these states. Such a potential neurobiological mechanism also aligns with the recently proposed Epileptor model of seizure dynamics, where any brain network might be capable of seizure generation depending on its vulnerability to crossing a critical separatrix barrier [36, 37]. In the framework of the Epileptor, our results suggest that synchronizing and desynchronizing nodes might regulate a critical level of network synchronizability and prevent the extent to which the network crosses a separatrix.

### 3.3 Relevance to Basic and Translational Neuroscience

We speculate that dysfunction of neural circuits in other brain network disorders might also be explained by irregularities in cortical push-pull control mechanisms. For instance, in healthy brain the excitatory and inhibitory pathways of the basal ganglia operate in concert as a push-pull system to control neural activity in the neocortex and brainstem for executive motor function [38, 39]. Imbalances in either of these pathways may lead to hypo-or hyperkinetic dysfunction of the basal ganglia in Parkinson,s Disease. The tools we present may provide a convenient way to vet new circuit targets in this disorder, when applied to multi-unit or local field recordings across these networks. Surgical lesioning of direct or indirect pathways via deep brain stimulation is a common method for re-balancing the putative push-pull control system and treating Parkinson,s. It is intuitively plausible that virtual cortical resection of the basal ganglia network may provide an opportunity to improve localization of control hubs as targets for stimulation.

Imbalances in synchronization have also been found in separate studies of patients with schizophrenia, autism, and Alzheimer,s disease [39]. Brain networks in schizophrenia may be unable to achieve a sufficient degree of synchronization between distant brain networks through corticothalamocortico loops. As a result of decreased control in local brain regions, significant local synchronization may underlie common symptoms such as hallucinations. Similar mechanisms of imbalances that result in hyperexcitation in autism and reduction of synchronization in Alzheimer,s disease have been suggested [39]. The virtual cortical resection method we describe here might point to important network control regions as the source of dysfunction in adequate regulation of network synchronization.

While our results support push-pull mechanisms in meso-scale cortical networks, competitive binding strategies that the brain employs for internally coordinating dynamics in distributed networks through synchronization and desynchronization bear striking similarity to push-pull control mechanisms [40]. Push-pull relationships have also explained cognitive control in large-scale brain networks. Antagonistic interactions between internally oriented processing in default mode network and a more distributed external attention system may support the notion that push-pull control assists in dynamically balancing metabolic resources geared towards introspective and extrospective processing [41]. And, evidence for an error correction system that mediates goal-directed tasks has been found in network interactions between prefrontal brain regions
[42]. Further work has posited that the flexibility of connections that drives network reorganization may also explain how the brain regulates the degree of competition between opposing cognitive systems [43]. From the perspective of push-pull control, network flexibility might help titrate synchronizability within a critical boundary of order and disorder in distributed networks [7, 44]. We believe an in-depth exploration of virtual resection applied to brain networks during cognitive tasks, may pinpoint brain areas that coordinate widespread interactions in the brain via push-pull control strategies.

In more basic science applications, our methods could have a role in decomposing network interactions in a variety of circuit investigations. One important feature of our methods is that they can be applied across scales and modalities, as network measures can be applied to signals as diverse as videos from optogenetic recordings, to “inscopix” videos of large numbers of neuronal calcium images, to more standard multielectrode array studies distributed spatially across the brain. Applying our virtual resection technique across modalities could provide a unique approach to characterize circuit behavior across multiple scales and its components without a priori knowledge of functional divisions.

### 3.4 Methodological Considerations

An important clinical consideration related to this work is the sampling error inherent in any intracranial implantation procedure. Any of the techniques used to map epileptic brain usually yield incomplete representations of the epileptic network. It is not possible to fully record from the entirety of cortex in affected patients, at least not with technologies currently available. In some cases this might mean that neither seizure onset zones nor all regions of seizure spread are fully delineated. These are problems that are limitations of epilepsy surgery, though usually clinical symptoms, brain imaging and video recording of seizures steer recordings to reasonably target epileptic networks. If successful, at least for superficial neocortical networks, magnetoencephalography (MEG) studies may provide a non-invasive method to delineate epileptic networks without invasive electrode placement. Even with some degree of sampling error, in this work we apply virtual cortical resection to study a single topological metric: network synchronizability. This will guide recordings at least to regions coupled to networks, in worst case, and we expect to rapidly understand signature of main seizure generators that will help guide us to identify and interpret undersampled studies.

### 3.5 Clinical Impact

Isolating the natural control mechanisms of brain function is critical for clinical translation. Enhancing and disrupting these natural control mechanisms could be a viable approach for introducing therapy with implantable devices for network disorders like epilepsy, in addition to resective surgery. Current methods of treating drug-resistant epilepsy rely on surgical resection or, more recently, implantable devices. However, predicting network response to therapy remains challenging. The virtual cortical resection technique is a novel, objective method of probing robustness and fragility upon removing components of the epileptic network. Using this method, we pinpointed putative network controllers that may be crucial for seizure evolution - suggesting that resection of these regions may compromise key mechanisms to contain seizure activity.

This technique will require careful retrospective, and then prospective, trials to validate its utility. Critical challenges include: (i) how to target functional connections based upon removal of cortical tissue? And, (ii) to determine if network models can account for neural plasticity after node removal, such as unmasking of
latent cortical connectivity ([45])? By honing network models to better capture structural and functional relationships in the brain, virtual cortical resection may allow clinicians to predict response to therapy and provide a quantitative guide to what is now a process guided by manual interpretation of ECoG recordings.

Finally, the clinical implications of this technique may reach beyond guided electrode placement for antiepileptic devices. In particular, these studies might open the way towards more accurate electrophysiologically-guided cortical resection or perhaps pinpointed thermal ablation to specific network regions, similar to procedures performed by cardiac electrophysiologists. These potential applications, while not immediately possible, offer considerable clinical advantages over the large cortical resections performed currently, with modest seizure-freedom rates. It is also likely that they will open the door to a host of other applications in movement, cognitive, affective and psychiatric disorders.

## 4 Methods

### 4.1 Patient Data Sets

#### 4.1.1 Ethics Statement

All patients included in this study gave written informed consent in accordance with the Institutional Review Board of the University of Pennsylvania.

#### 4.1.2 Electrophysiology Recordings

Ten patients undergoing surgical treatment for medically refractory epilepsy believed to be of neocortical origin underwent implantation of subdural electrodes to localize the seizure onset zone after presurgical evaluation with scalp EEG recording of ictal epochs, MRI, PET and neuropsychological testing suggested that focal cortical resection may be a therapeutic option. Patients were then deemed candidates for implantation of intracranial electrodes to better define epileptic networks. De-identified patient data was retrieved from the online International Epilepsy Electrophysiology Portal (IEEG Portal) [46].

ECoG signals were recorded and digitized at 500 Hz sampling rate using Nicolet C64 amplifiers and pre-processed to eliminate line noise. Cortical surface electrode (Ad Tech Medical Instruments, Racine, WI) configurations, determined by a multidisciplinary team of neurologists and neurosurgeons, consisted of linear and two-dimensional arrays (2.3 mm diameter with 10 mm inter-contact spacing) and sampled the neocortex for epileptic foci (depth electrodes were first verified as being outside the seizure onset zone and subsequently discarded from this analysis). Signals were recorded using a referential montage with the reference electrode, chosen by the clinical team, distant to the site of seizure onset and spanned the duration of a patient,s stay in the epilepsy monitoring unit. See Table 1 for demographic and clinical information.

**Table 1:** Patient information. Patient data sets accessed through IEEG Portal (http://www.ieeg.org). Age at first reported onset and at phase II monitoring. Localization of seizure onset and etiology is clinically-determined through medical history, imaging, and long-term invasive monitoring. Seizure types are SP (simple-partial), CP (complex-partial), CP+GTC (complex-partial with secondary generalization). Counted seizures were recorded in the epilepsy monitoring unit. Clinical imaging analysis concludes L, Lesion; NL, non-lesion. Surgical outcome was based on Engel score (scale: I-IV, seizure freedom to no improvement; NR, no-resection; NF, no follow-up). M, male; F, female.

| Patient<br>(IEEG Portal) | Sex | Age (Years)<br>(Onset/Surgery) | Seizure Onset | Etiology | Seizure Type<br>(N) | Imaging | Outcome |
| --- | --- | --- | --- | --- | --- | --- | --- |
| HUP64_phaseII | M | 03/20 | Left frontal | Dysplasia | CP+GTC (1) | L | I |
| HUP65_phaseII | M | 02/36 | Right temporal | Auditory reflex | CP+GTC (3) | N/A | I |
| HUP68_phaseII | F | 15/26 | Right temporal | Meningitis | CP (1),<br>CP+GTC (4) | NL | I |
| HUP70_phaseII | M | 10/32 | Left<br>perirolandic | Cryptogenic | SP (8) | L | NR |
| HUP72_phaseII | F | 11/27 | Bilateral left | Mesial temporal<br>sclerosis | CP+GTC (1) | L | NR |
| HUP73_phaseII | M | 11/39 | Anterior right<br>frontal | Meningitis | CP+GTC (5) | NL | I |
| HUP78_phaseII | M | 00/54 | Anterior left<br>temporal | Traumatic<br>injury | CP (5) | L | III |
| HUP79_phaseII | F | 11/39 | Occipital | Meningitis | CP (2) | L | NR |
| HUP86_phaseII | F | 18/25 | Left temporal | Cryptogenic | CP+GTC (2) | NL | II |
| HUP87_phaseII | M | 21/24 | Frontal | Meningitis | CP (2) | L | I |

#### 4.1.3 Description of Epileptic Events

We analyzed 18 partial seizures (simple and complex) and 16 partial seizures that generalized to surrounding tissue, forming a population of focal and distributed epileptic networks, respectively. Seizure type, onset time, and onset localization were marked as a part of routine clinical workup.

The seizure state spanned the period between clinically-marked earliest electrographic change (EEC) [28] and termination; and the pre-seizure state spanned a period equal in duration to the seizure state and ended immediately prior to the EEC. Note that we refer to each pair of pre-seizure and seizure states as an *event*.

### 4.2 Functional Network Construction

#### 4.2.1 Pre-Processing

ECoG signals from each event were divided into 1s, non-overlapping, weakly stationary time-windows in accord with related studies [16]. In each time window, signals were re-referenced to the common average reference [16, 47] to account for variation in reference location across patients and to avoid broad field effects that may bias connectivity measurements erroneously in the positive direction.

#### 4.2.2 Coherence Estimation

We constructed functional networks in each time-window using multitaper coherence estimation, which defines a network connection between electrode pairs as the power spectral similarity of signal activity over a specific frequency band. We applied the *mtspec* Python implementation [48] of multitaper coherence estimation with time-bandwidth product of 5 and 8 tapers in accord with related studies [49]. Based on a vast literature implicating high-frequency oscillations and γ activity as drivers of epileptic activity, we primarily studied functional connectivity in the high-γ band (95–105 Hz). This frequency range represents relatively local neural population dynamics that are largely unaffected by volume conduction. See the Supplement for complementary results in the lower γ band.

### 4.3 Metrics of the Time-Varying Functional Network

#### 4.3.1 Network Geometry

In our network analysis, we refer to heterogeneity of network architecture in the context of node strength, also known as weighted degree. We measure a time-varying quantity of heterogeneity: the dispersion of the degree distribution in each time window. We use a non-parametric, interquartile distance measure to quantify dispersion. The interquartile distance is the 75th percentile subtracted by the 25th percentile of a distribution.

#### 4.3.2 Network Synchronizability

A recent trend in studying functional networks is to model dynamic geometric structure that evolves through system states [12, 9, 10, 11]. Building on the classical notion of stability of the synchronized state in static networks, we fit a popular synchronizability model for dynamic networks [50] to account for time-varying structure of the functional networks in our study. As a simplification for our analysis, we analyze each time-window separately.

To quantify synchronizability, we first estimated the time-varying Laplacian matrix **L**(*t*) for each time-window *t* of the functional network. Intuitively, each entry *l_ij_*(*t*) of **L**(*t*) can be interpreted as a measure of how easily information could diffuse between nodes i and j based on the relative connectivities of both nodes to all other nodes in the network. Next, we computed the eigenspectrum of **L**(*t*) and calculated the ratio of the second-smallest eigenvalue λ_2_ to the largest eigenvalue λ_max_ for each *t*, resulting in network synchronizability 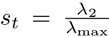(larger values of *s(t)* correspond to greater state stability) [29]. The **Supplementary Note** provides details of the *master stability function* formalism behind synchronizability and its relationship to state stability.

#### 4.3.3 Virtual Cortical Resection

To model potential effects of resecting or lesioning regions of brain networks, we develop a virtual cortical resection technique. Generally, the approach allows us to ask how network topology might change upon removing one or more nodes or connections in the network. In time-varying networks, virtual cortical resections may be useful in patterned lesioning schemes for implantable devices that continuously modulate brain state away from seizures [5, 6].

Here, we tailored virtual cortical resection to study putative controllers that regulate synchronizability in the epileptic network. We measure the control centrality, the contribution of a node to network synchronizability, by applying virtual cortical resection to each node in each time window of the functional network. The control centrality identifies a node as a desynchronizing (c_i_ > 0) or synchronizing (c_i_ < 0) controller. The Supplementary Material explores the relationship between control centrality and other topological properties of networks.

#### 4.3.4 Statistical Analysis

We compared time-varying network metrics between partial seizures that remain focal and partial seizures that generalize to surrounding tissue. We performed this comparison by (i) normalizing each seizure event into 20 sequential time-bins spanning the pre-seizure and seizure states and (ii) employing functional data analysis to statistically test differences in temporal dynamics between seizure type, independently in each state. We assigned p-values to each state by re-assigning events uniformly at random to seizure types up to 1,000,000 times and computing the mean area between the resulting functional curves.

## 5 Acknowledgments

AK and BL acknowledge support from the National Institutes of Health through awards #R01–NS063039, #1U24 NS 63930–01A1, Neil and Barbara Smit, the Citizens United for Research in Epilepsy (CURE) through Julie,s Hope Award, and the Mirowski Foundation. DSB acknowledges support from the John D. and Catherine T. MacArthur Foundation, the Alfred P. Sloan Foundation, the Army Research Laboratory and the Army Research Office through contract numbers W911NF–10–2–0022 and W911NF–14–1–0679, the National Institute of Child Health and Human Development (1R01HD086888–01), the Office of Naval Research, and the National Science Foundation (BCS-1441502 and BCS-1430087). The funders had no role in study design, data collection and analysis, decision to publish, or preparation of the manuscript.

